# Small extracellular vesicles mediate the antihyperalgesic effect of bone marrow stromal cells: the role of “priming”

**DOI:** 10.64898/2026.05.08.723785

**Authors:** Wei Guo, Jia-Le Yang, Huakun Xu, Kamal D. Moudgil, Feng Wei, Ke Ren

## Abstract

Multipotent mesenchymal stem cells (MSCs) including bone marrow stromal cells (BMSCs) have shown analgesic efficacy in recent years. Studies suggested that the therapeutic effect of MSCs was mediated by their secreted small extracellular vesicles (sEVs) mainly exosomes. The present study evaluated the antihyperalgesic effect of BMSC-related sEVs in a mouse model of neuropathic pain involving chronic constriction injury of the infraorbital nerve (CCI-ION). Our separation protocol generated EV particles mostly sized in the range of exosomes (30-170 nm) and express exosome marker proteins CD9, CD81, and Tsg101, suggesting their endosome origin.

We show that intravenous injection of BMSC-related sEVs attenuated pain hypersensitivity induced by CCI-ION as indicated by decreased mechanical hypersensitivity (von Frey test) and reduced aversion to noxious stimulation (conditioned place avoidance test). The antihyperalgesic effect of sEVs was observed in both female and male animals, and the effect was dose-dependent. sEVs from *NAIVE* serum-treated BMSC cultures produced short-lasting antihyperalgesia in male but not female mice, suggesting a subtle sex difference. The antihyperalgesia of sEVs from BMSC culture was blocked by the pretreatment of the culture with GM4869, the antagonist of exosome secretion, suggesting that the effect was not related to other co-isolated soluble mediators but mediated by MSC-derived exosomes.

Interestingly, the *prior injury condition* in which sEVs were isolated favors the pain-relieving effect of sEVs. sEVs isolated from the serum of BMSC-treated animals receiving tendon ligation (TL) injury attenuated hyperalgesia for 24 h, while sEVs from the serum of BMSC-treated *NAIVE* animals only attenuated hyperalgesia at 3 h after injection. sEVs from the BMSC culture treated with the serum of TL rats were antihyperalgesic, but sEVs from the BMSC culture treated with the serum of naive animals were ineffective.

Our results indicate that BMSC-related sEVs produced antihyperalgesia similar to that produced by BMSCs. The results suggest that the interactions between BMSCs and injury conditions are crucially important for producing efficacious sEVs/exosomes and support that the effect of sEVs could be optimized by priming BMSCs with injury-related conditions.

## Introduction

Chronic pain is a debilitating condition that affects tens of millions of people in the US (Dahlhamer et al., 2018). The current treatment for chronic pain, such as neuropathic and musculoskeletal pain, is unsatisfactory. Facing the current opioid epidemic, the need for alternative pain management has become even more pressing. One promising alternative pain management approach is to use cell-based therapy. Multipotent mesenchymal stem cells (MSCs), including bone marrow stromal cells (BMSCs), have shown analgesic efficacy in preclinical studies (Siniscalco et al., 2010; Guo et al., 2011,2017; Chen et al., 2015). Clinical studies have also shown the pain-reducing effects of MSCs (Black et al., 2007; Gupta et al., 2013; Guo et al., 2014; Vickers et al., 2014; Vaquero et al., 2018; Ren et al., 2023). Studies on the therapeutic mechanisms of MSCs identified that the effect of MSCs was mediated by their secreted exosomes, or small extracellular vesicles (sEVs) (Lai et al., 2010).

Exosomes are a class of sEVs secreted by all types of cells (see Ren 2019, for a review). The literature suggests that the therapeutic outcome of MSCs could be mimicked by a concentrated MSC-conditioned medium that contains a secretome of MSCs (Timmers et al., 2007; Gama et al., 2018) and that the therapeutic effect of MSCs is attributable to their exosomes (Casado et al., 2017; Gonzalez-King et al., 2017; Kou et al., 2018). Kordelas et al. (2014) showed that MSC-derived exosomes significantly improved the symptoms of graft-vs-host disease, suggesting that MSC-exosomes are a promising surrogate of MSC-based therapy. Further, recent preclinical studies have shown the pain-relieving effect of MSC-derived exosomes or sEVs. Intrathecal injection of MSC exosomes reduced neuropathic pain in rats (Shiue et al., 2019). MSC-derived exosomes attenuated persistent pain in animal models of osteoarthritis (Zhang et al., 2019; He et al., 2020; Li et al., 2020). Extracellular vesicles from MSCs attenuated chronic pelvic pain in a rat autoimmune prostatitis model (Peng et al., 2021).

This study was undertaken to seek to redefine and improve the use of therapeutic MSCs in pain relief. We evaluated the antihyperalgesic effect of BMSC-related sEVs in a mouse model of neuropathic pain involving chronic constriction injury of the infraorbital nerve (CCI-ION) (Wei et al., 2008; Okubo et al., 2013). We further tested a hypothesis that the interactions between BMSCs and injury conditions are crucially important for producing efficacious sEVs and that the effect of sEVs could be optimized by priming BMSCs with injury-related conditions.

## METHODS

### Animals

Adult female and male C57Bl/6 mice were purchased (Envigo-Harlan, Frederick, MD). Female adult Sprague Dawley rats (Envigo-Harlan) were used as donors of BMSCs and blood. We used rat BMSCs in the present study since, compared to the mouse, the yield of BMSCs is higher in the rat, and fewer donor animals were needed for the study. Rat BMSCs worked well in mice, and mice have well tolerated the rat BMSCs (Guo et al., 2017). Animals were kept under controlled environmental conditions (≈22°C), relative humidity 40-60%, 12h/12h light-dark cycles, and food and water *ad libitum*. All surgical procedures were performed under Ketamine (100-150 mg/kg)/Xylazine (10-16 mg/kg) anesthesia (i.p.). The CCI-ION mouse model was produced via an intraoral approach according to Wei et al. (2008). A 5-7 mm long incision was made along the gingivobuccal margin in the buccal mucosa. The ION was freed from surrounding connective tissues. At 3-4 mm from the nerve where its branches emerge from the infraorbital fissure, the ION was loosely tied with two chromic gut (4.0) ligatures, 2 mm apart. The masseter muscle tendon ligation mouse model (TL) was produced via a similar procedure except that the tendon of the masseter muscle was freed and ligated after the gingivobucal incision (Guo et al., 2010). All experiments were carried out following the National Institute of Health Guide for the Care and Use of Laboratory Animals (NIH Publications No. 80-23) and approved by the University of Maryland, Baltimore, Institutional Animal Care and Use Committee, School of Medicine.

### Measurement of mechanical sensitivity

The mechanical sensitivity test of the orofacial region was adapted from rats (Guo et al., 2010, 2019). The mouse was habituated to be held entirely in the experimenter’s hand, wearing a leather work glove. A series of calibrated von Frey filaments was applied to the skin within the infraorbital territory, near the center of the vibrissa pad on hairy skin surrounding the mystacial vibrissae or above the injured muscle. An active withdrawal of the head from the probing filament was defined as a response. Each von Frey filament was applied 5 times at intervals of 5-10 seconds. The response frequencies (the number of responses/number of stimuli) X100%) to a range of von Frey filament forces were determined, and a stimulus-response frequency (S-R) curve was plotted. After non-linear regression analysis, an EF_50_ value, defined as the effective von Frey filament force (g) that produces a 50% response frequency, was derived from the S-R curve (Prism, GraphPad) (Guo et al., 2004). CCI/ION and TL led to a leftward shift of the S-R curve and a reduction of EF_50_, suggesting the development of mechanical hypersensitivity, or hyperalgesia/allodynia.

### Measurement of conditioned place avoidance behavior (CPA)

For the CPA test (LaBuda & Fuchs 2000, Guo et al., 2019), animals were placed within a 42 (L) x 18 (W) x 18 (H) cm Plexiglas chamber. One half of the chamber was made of white plates (light area) and the other half was by black plates (dark area). During the test, animals were allowed unrestricted movement throughout the test chamber for the 30-min test period. Mice normally prefer to stay in dark areas. Testing began immediately with suprathreshold mechanical stimulation (11 g von Frey monofilament) applied to the facial skin at 15-sec intervals. The mechanical stimulus was applied to the skin above the injured nerve or muscle when the animal was within the preferred dark area and to the facial skin on the non-injured side when the animal was within the non-preferred light side. Based on the location of the animal at each 15-sec interval, the mean percentage of time spent in the black (dark) or white (light) chamber was calculated. Since staying in the preferred dark area was associated with an aversive and painful stimulus after CCI or TL, the injured mouse tended to leave the dark side to spend time on the unpreferred white side. An increase in the time spent in the light area suggests an increase in stimulus aversiveness.

### BMSC procedures

BMSCs were obtained from rats as described (Guo et al., 2011). The rats were sacrificed with CO_2_, and both ends of the tibiae, femurs, and humeri were cut off by scissors. A syringe fitted with an 18-gauge needle was inserted into the shaft of the bone, and bone marrow was flushed out with culture medium (alpha–modified Eagle medium, Gibco, Carlsbad, CA, USA; 10% fetal bovine serum (FBS), Hyclone, Logan, UT, USA). The bone marrow was then mechanically dissociated, and the suspension was passed through a 100-μm cell strainer to remove debris. The cells were incubated at 37°C in 5% CO_2_ in tissue-culture flasks (100 X 200 mm) (Sarstedt, Nümbrecht, Germany), and non-adherent cells were removed by replacing the medium. The property of expanded cells was assessed by flow cytometry with conventional markers (Guo et al., 2011).

On day 7, when the cultures reached 80% confluence, the cells were washed with PBS and harvested by incubation with 1 ml of 0.25% trypsin/1 mM ethylenediaminetetraacetic acid for 2 min at room temperature. Trypsin was neutralized by adding 5 ml of the complete medium. Cells were centrifuged at 1,000 X g for 2 min, the supernatant was removed, and the pellet was washed with PBS. The cell numbers were calculated by the Hemacytometer. For intravenous administration, 1.5 × 10^^6^ cells in 0.2 ml PBS were slowly injected into one tail vein of the anesthetized rat over 2 minutes using a 22-gauge needle.

### Small extracellular vesicles/exosomes procedures

For serum sEV separation, blood was collected from rats, placed into BD serum collecting tubes, and kept at room temperature for one hour. The blood was centrifuged at 2,000 X g for 30 min at 4°C. The serum was collected from the supernatant and filtered through a 0.22 μm filter. One ml of serum was used for isolating the exosome using Invitrogen Total Exosome Precipitation Reagent (from serum, Cat. #4478360) per the manufacturer’s instructions. For BMSC sEVs, one week after seeding, the culture medium was replaced with an FBS-free medium (MEM Alpha, Gibco). The BMSC culture (1 × 10^^6^ cells) was treated with serum (20%, 37°C, 16 hr) from naive rats or rats with a tendon injury. The culture medium was collected, centrifuged at 2000g for 30 min, and filtered. Ten ml medium was used to isolate exosomes by using the Total Exosome Isolation Kit for cell culture media (Invitrogen, Cat. #4478359). sEV particles were evaluated by electron microscopy and flow cytometry (UMB Cores), and nanoparticle tracking analysis with Particle Metrix’ ZetaView (AlphaNanotech, NC) (Fig. 1) (Bachurski et al 2019). Total sEV protein concentrations were determined (BCS protein assay kit, Pierce).

**Fig. 1.**
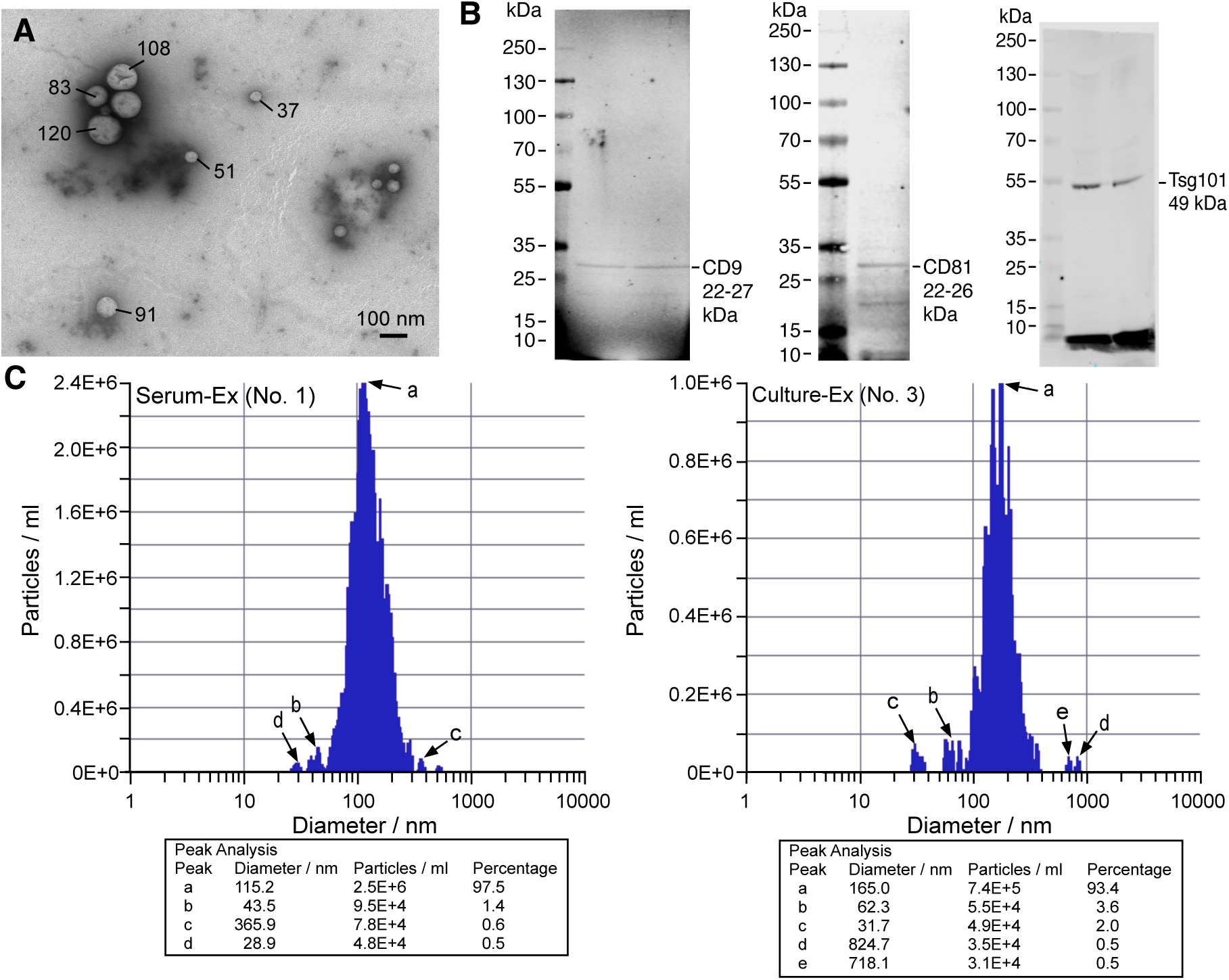
Characterization of sEVs/exosomes derived from naïve rats. **A**. sEVs were isolated, and samples were applied to glow-discharged 400 mesh formvar-coated copper grids, negatively stained with 1% (wt/vol) uranyl acetate, air-dried, and examined in a transmission electron microscope (Tecnai T12, Thermo Scientific) at an operating voltage of 80 kV. Digital images were acquired using an AMT bottom-mount CCD camera and AMT600 software. The numbers indicate the diameters of EVs in nm. **B**. Western blot illustrating exosome-enriched protein markers CD9, CD81, and Tsg101. The CD9 and CD81 samples were from the serum and the Tsg101 samples were from the culture medium of BMSCs. **C**. Size distribution of exosome particles. Sample No. 1 was from the serum (left), and sample No. 3 (right) was from the culture medium. Peak analysis was performed on selected peaks (a-e).

### Western blot

Proteins of separated sEVs were isolated by Total Exosome RNA & Protein Isolation kit per the manufacturer’s protocol (Invitrogen). The protein concentration was determined using a detergent-compatible protein assay with a bovine serum albumin standard. For detecting the immunoreactivity with near-infrared fluorescence using the Odyssey Infrared Imaging System (Odyssey®CLx, LI-COR, Lincoln, NE), 50 μg protein samples were denatured by boiling for 5 min and loaded onto 4-20% Bis-Tris gels (Invitrogen). After electrophoresis, proteins were transferred to nitrocellulose membranes. The membranes were blocked for 1 h with Odyssey Blocking Buffer and then incubated with primary antibodies diluted in Odyssey Blocking Buffer at 4^°^C overnight, followed by washing with PBS containing 0.1% Tween 20 (PBST) three times. The membranes were then incubated for 1 h with IRDye800CW-conjugated goat anti-rabbit IgG and IRDye680-conjugated goat anti-mouse IgG secondary antibodies (LI-COR) diluted in Odyssey Blocking Buffer. The blots were further washed three times with PBST and rinsed with PBS. Proteins were visualized by scanning the membrane with 700- and 800-nm channels. The loading and blotting of the amount of protein were verified by re-probing the membrane with anti-β-actin antibodies.

### Drugs and antibodies

GW4869 (Sigma-Aldrich), a cell-permeable, symmetrical dihydroimidazolo-amide compound that acts as an inhibitor of neutral sphingomyelinase and is widely used as an exosome/sEV inhibitor (Menck et al., 2017), was dissolved in 5% dimethyl sulfoxide (DMSO) and saline. The following rabbit monoclonal antibodies used in western blot were from Abcam: anti-CD9 (1:500, ab223052), anti-CD81 (1:500, ab155760), and anti-TSG101 (1:1000, ab133586). PE anti-mouse/rat CD81 antibody (Cat. No. 104905) from Biolegend was used in flow cytometry. Mouse monoclonal anti-β-actin antibodies were from Sigma-Aldrich (1:5,000).

### Data analysis

All data collection/analyses were conducted under blind conditions whenever possible. Power analyses were performed to determine the number of animals needed to provide an 80% chance to achieve statistical significance (alpha = 0.05). Data are presented as mean ± S.E.M. Statistical comparisons were made by ANOVA followed by *post hoc* comparisons with Bonferroni corrections. For animals that were subject to repeated testing, ANOVA with repeated measures was used. P < 0.05 was considered significant for all cases.

## RESULTS

The separated sEVs appeared round, mostly with a diameter between 30-170 nm and positive for tetraspanins CD9 and CD81, and other exosome markers such as Tsg101 (Fig. 1). ZetaView particle analysis showed a predominant size distribution peak at 115 nm for serum-derived (“a” in Fig. 1C, left) and 165 nm for BMSC culture-derived (“a” in Fig. 1C, right) particles, respectively. The characteristics of the isolated sEVs suggest their exosome identity.

### sEVs derived from BMSC-treated animals mimic the pain-relieving effect of BMSCs

Our first goal was to determine whether exosomes derived from animals receiving transplantation of BMSCs produced the pain-relieving effect. Systemic injection of BMSCs attenuates behavioral hyperalgesia (Guo et al., 2011). To mimic this effect, we isolated sEVs/exosomes from the serum of the TL (BMSC-TL) or naive (BMSC-Naive) rats receiving BMSCs (1 X 10^6^ cells/0.2 ml), or TL rats receiving BMSC culture medium (Med-TL) (Fig. 2A). CCI-ION was produced in female mice, which led to long-lasting pain hypersensitivity, or hyperalgesia, as shown by a significant and persistent reduction of EF_50_s in the affected orofacial region (Fig. 2B). Injection of sEVs (0.02 ml, i.v.) derived from BMSC-treated TL animals led to an increase in EF_50_s at 3 h and 1 d after injection, indicating attenuation of CCI-induced hyperalgesia, compared to the CCI-4w time point (p<0.01-0.001) and sEVs from culture medium-treated TL animals (Med-TL, p<0.001) (Fig. 2B). BMSC-Naive sEVs attenuated hyperalgesia only at 3 h after injection (p<0.05). BMSC-TL sEVs also reduced CCI-produced aversion as shown by a CPA test. At 1 d after injection of sEVs, the CCI-mice receiving BMSC-TL sEVs spent significantly less amount of time in the light chamber as compared to animals receiving Med-TL or BMSC-Naive sEVs (p<0.001) (Fig. 2C). Similarly, BMSC-TL sEVs also produced pain attenuation in the TL mice (Supplemental Fig. 1). Fig. 3 shows that the effect of BMSC-TL sEVs was dose-dependent. Consistent with Fig. 2B, undiluted BMSC-TL sEVs attenuated hyperalgesia at 3 h-1 d after injection. A 10-fold dilution (0.1x) of the sEVs only produced pain attenuation at 3h, and a 100-fold (0.01x) diluted sEVe solution produced a slight increase in EF_50_s at 1 d after injection (Fig. 3). Thus, BMSC-related sEVs produced antihyperalgesia similar to that of BMSCs, and importantly, prior interactions between BMSCs and injury conditions favored the therapeutic effect of sEVs.

**Fig. 2.**
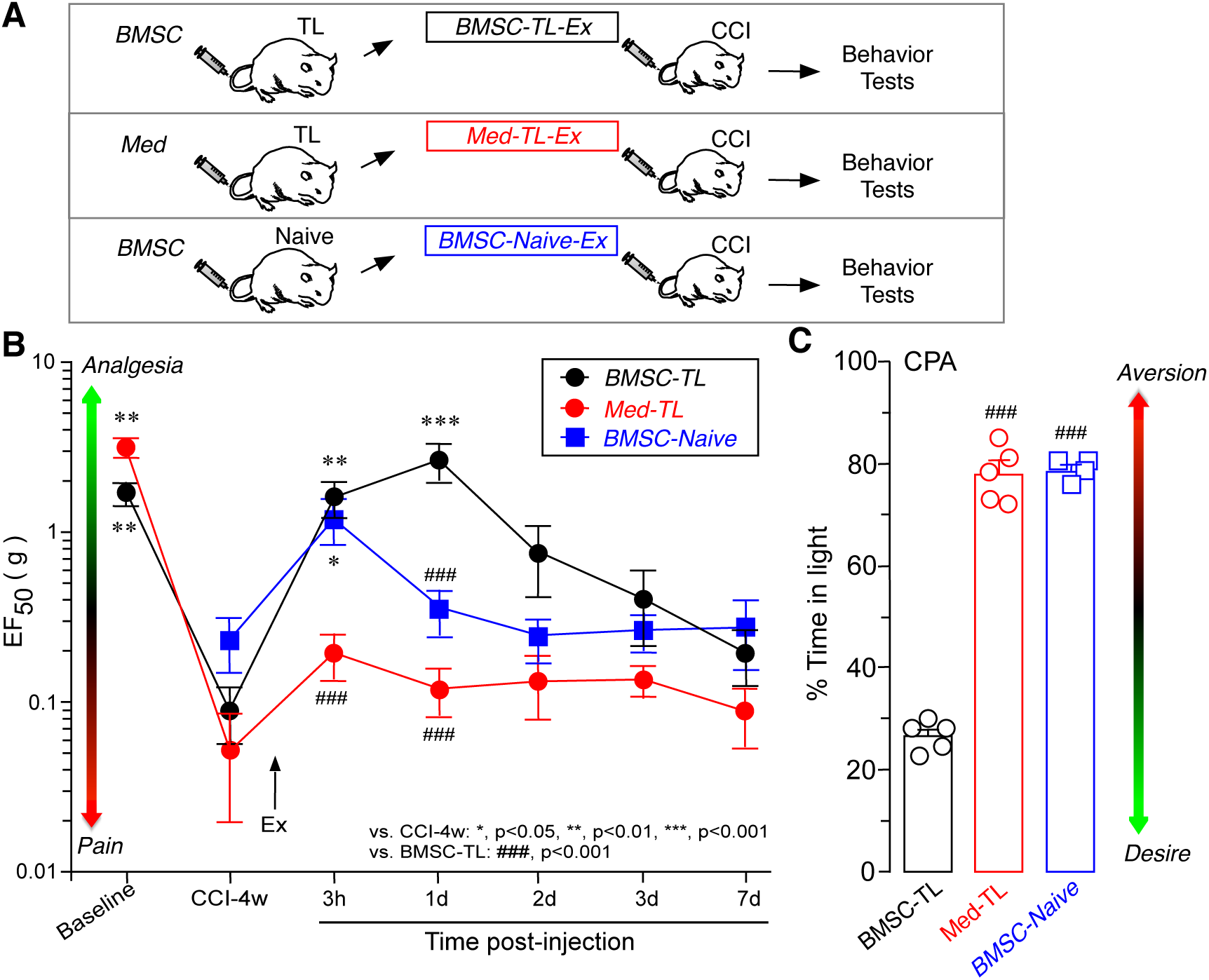
Anti-hyperalgesia by serum-derived sEVs/exosomes. **A**. Flowchart of the experiment. BMSCs or culture medium was injected (i.v.) into the TL or naive rats, and exosomes were isolated from the serum of the donor animals. Isolated exosomes were injected into mice at 4 w after CCI, and behavioral tests were conducted at 3 h to 7 d after exosome injection. **B**. An IV injection of exosomes (0.02 ml) isolated from BMSC-treated TL animals (BMSC-TL) elevated EF_50_ at the 3 h and 1 d timepoints, indicating attenuation of behavioral hyperalgesia in CCI mice as compared to exosomes from culture medium-treated animals (Med-TL). Exosomes from BMSC-treated naïve animals (BMSC-Naive) attenuated hyperalgesia at 3 h after injection. **C**. BMSC-TL exosome reduced % of Time in the light chamber in CCI mice at 1 d after injection. *, p<0.05, **, p<0.01, ***, p<0.001, vs. CCI-4w; ###, p<0.001, vs. BMSC-TL. N=4-5/Group. Post-hoc comparison after one-way or two-way ANOVA with repeated measures.

**Fig. 3.**
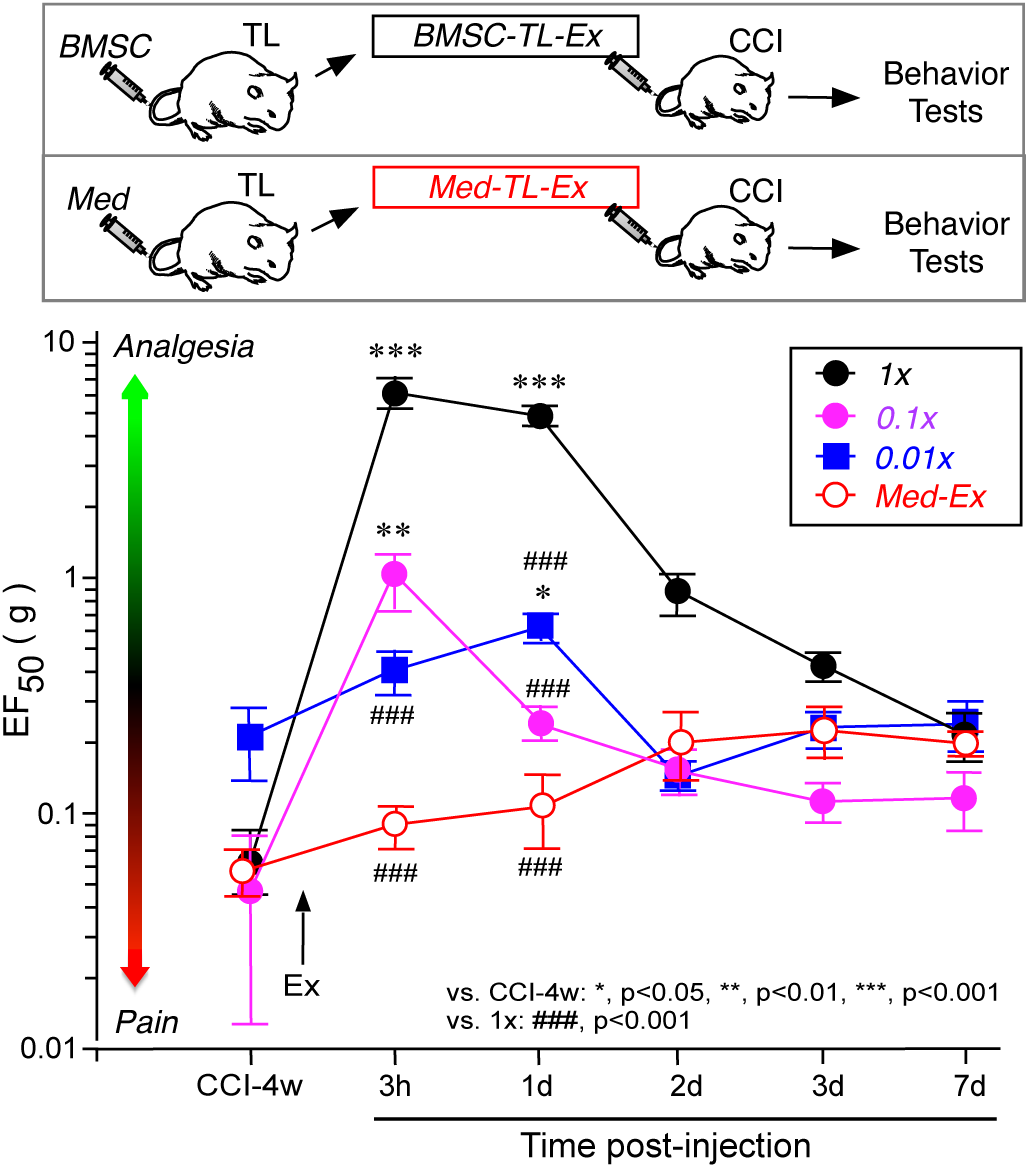
The dose dependency of sEVs/exosome-produced anti-hyperalgesia. The flow of the experiments is shown on top. sEVs/exosomes were isolated from the serum of TL rats receiving BMSCs (1.5x10^6^) and injected into mice at 4 weeks after CCI. The doses of BMSC-exosomes were 1x (undiluted), 0.1x (10-fold dilution), and 0.01x (100-fold dilution). Exosomes from the serum of TL rats receiving culture medium (Med-Ex) were used as controls. Note the dose-dependent effect of BMSC-Ex at 3 h and 1 d after injection. *, p<0.05, **, p<0.01, ***, p<0.001, vs. CCI-4w; ###, p<0.001, vs. BMSC-Ex at 1X. N=5/Group. Post-hoc comparisons after one-way or two-way ANOVA with repeated measures.

### The antihyperalgesic effect of sEVs directly derived from BMSC cultures

Exosomes from the serum are diverse and some of them contain mediators related to the development of pain hypersensitivity (Jean-Toussaint et al., 2020). We next separated sEVs directly from BMSC cultures and examined their effect on persistent pain. We also further compared the effect of “priming” on the antihyperalgesic effect of sEVs. One week after seeding, BMSC cultures (1 × 10^^6^ cells) were treated with serum from TL or naive rats for 16 h, and sEVs were separated from the culture medium (Fig. 4A). Injection of TL-BMSC culture-derived sEVs produced dose-dependent antihyperalgesia in female CCI-ION mice, which was evident at 3 h and 1 d after sEV injection (Fig. 4B). On the other hand, sEVs from BMSC cultures pretreated with naive serum did not attenuate pain. The CPA test showed reduced time in the light chamber in mice receiving sEVs from TL serum-pretreated cultures, compared to sEVs from naive serum-treated BMSC cultures (Fig. 4C). TL-BMSC sEVs also attenuated pain hypersensitivity in male mice (Supplemental Fig. 2).

**Fig. 4.**
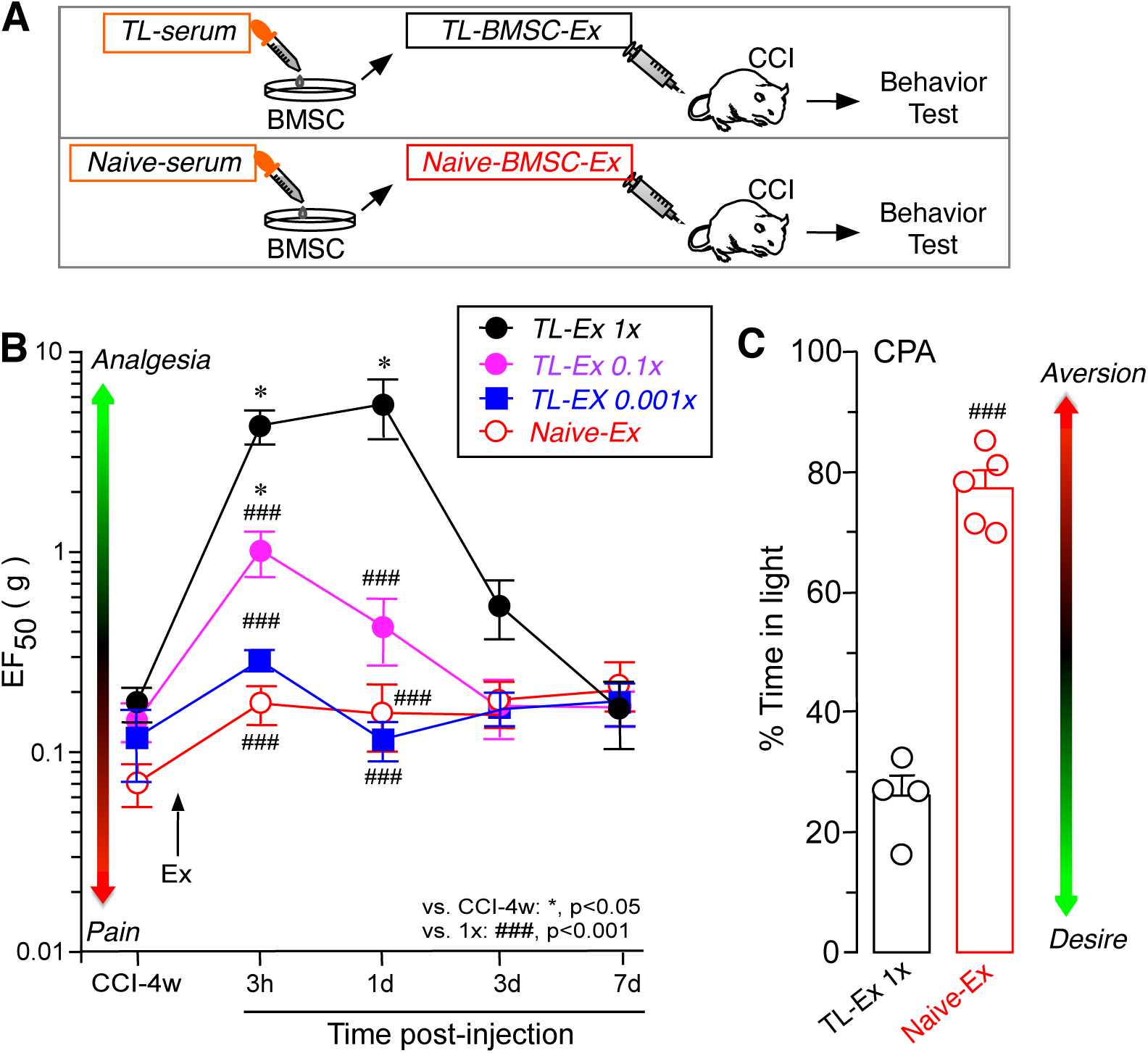
The effect of BMSC culture-derived sEVs/exosomes on behavioral hyperalgesia. **A**. Flowchart of the experiment. The BMSC culture was primed with the serum (20%, 37°C, 16 h) from TL or naive rats one week after seeding. Ten milliliters of culture medium were used to isolate exosomes. **B**. An IV injection of exosomes (0.02 ml) isolated from the BMSC culture primed with serum from TL animals (TL-Ex 1x) elevated EF_50_ in CCI mice, compared to exosomes from the BMSC culture treated with serum from naïve animals (Naïve-Ex). The effect of TL-Ex was dose-dependent. **C**. TL-Ex reduced % Time in the light chamber in CCI mice at 1 d after injection, suggesting a reduced aversion to a noxious stimulus. *, p<0.05, vs. CCI-4w; ###, p<0.001, vs. TL-Ex 1x. N=4-5/Group. Post-hoc comparisons after one-way or two-way ANOVA with repeated measures.

The use of culture-derived sEVs allowed us to test the effect of the sEV/exosome antagonist. GM4869 (10 μM) or vehicle was added to the BMSC culture two hours before treating with TL-serum, and sEVs were separated and injected into the CCI mice (Fig. 5A). Pretreatment of BMSC culture with GM4869 eliminated the antiyhyperalgesic effect of sEVs, while vehicle-treated sEVs produced pain attenuation and reduction of aversion consistent with above experiments (Fig. 5B, C). These results suggest that exosome secretion from BMSCs is required for their pain-relieving effect.

**Fig. 5.**
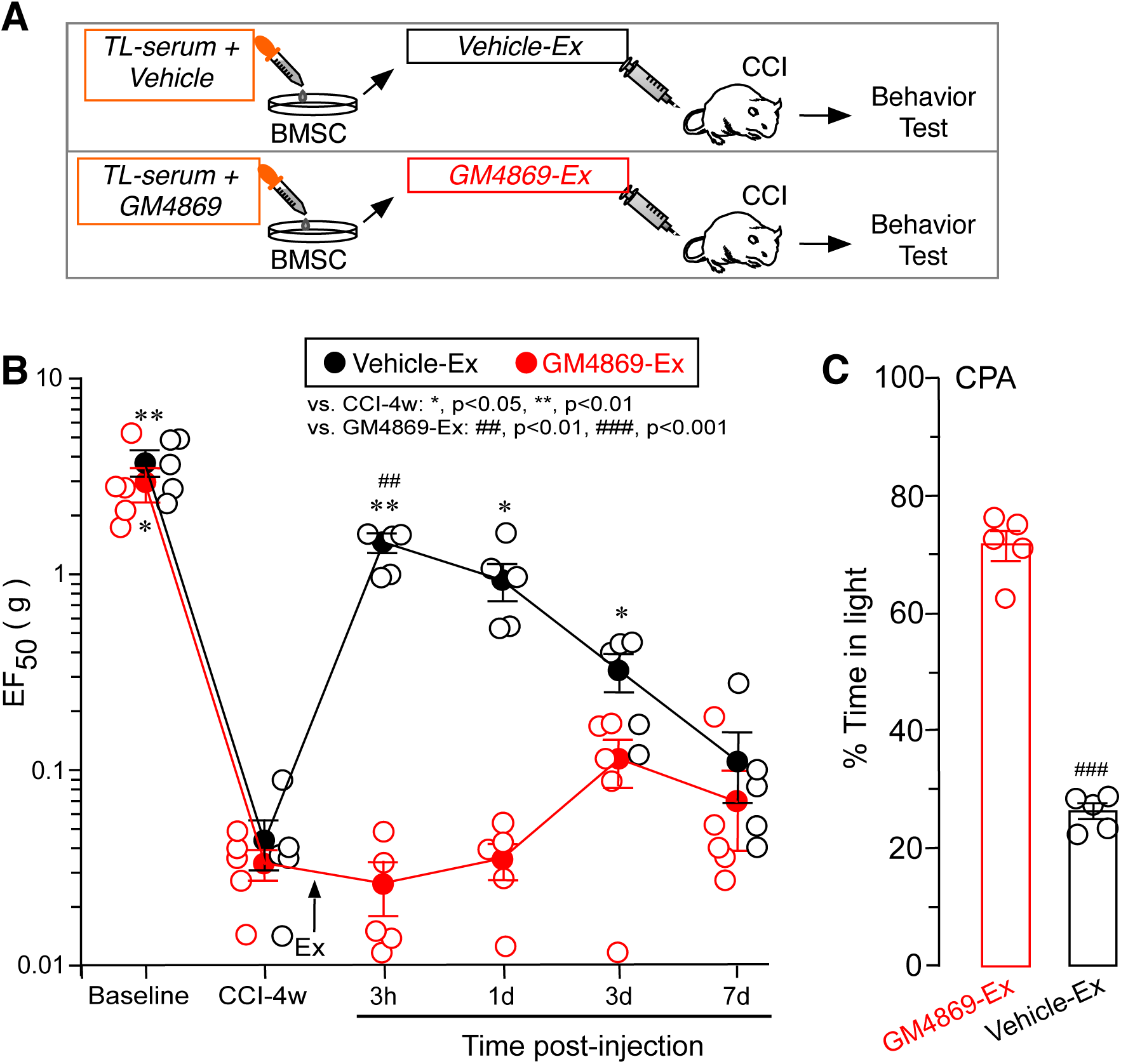
The effect of GM4869 on sEVs/exosome-produced antihyperalgesia. **A**. Flowchart of the experiment. The BMSC culture was pre-treated with the serum (20%, 37°C, 16 h) from TL rats one week after seeding. GM4869, an exosome secretion inhibitor or vehicle, was added to the culture with serum before exosome isolation. Ten milliliters of culture medium were used to isolate exosomes. **B**. Compared to exosomes from vehicle-treated culture (Vehicle-Ex), the addition of GM4869 to the culture reversed exosome-produced antihyperalgesia. **C**. GM4869-Ex-treated mice showed a higher preference for the light chamber, compared to Vehicle-Ex, suggesting an increased aversion to a noxious stimulus. *, p<0.05, **, p<0.01, vs. CCI-4w; ##, p<0.01, ###, p<0.001, vs. GM4869-Ex. N=5/Group. Post-hoc comparisons after one-way or two-way ANOVA with repeated measures.

## DISCUSSION

Extracellular vesicles consist of three major groups of nanoparticles: exosomes originating from the endosome compartment, ectosomes (microparticles) from the cell membrane, and apoptotic bodies from dying cells. Although each subclass of Evs differs in its features such as size, cargo, and membrane content, the overlap in size and molecular markers makes it practically difficult to achieve a pure preparation and establish their identity with confidence (Cocucci and Meldolesi, 2015; Witwer et al., 2019; Poupardin et al., 2021). The use of the generic term “small Extracellular Vesicles” in the present work reflects this fact. Currently, there are numerous EV isolation or separation protocols based on different techniques (Helwa et al., 2017; Poupardin et al., 2021). After preliminary testing of the traditional ultracentrifuge protocol and commercial kits, we chose the Invitrogen kit for our study as it produced consistent results with sufficient yield for our *in vivo* functional analyses. Our separation protocol generated EV particles mostly sized in the range of exosomes and expressed exosome marker proteins CD9, CD81, and Tsg101, suggesting their endosome origin. Functionally, exosomes and ectosomes may be analogous to each other (Cocucci and Meldolesi, 2015), but exosomes have been shown more efficacious clinically as they produce greater immune suppression in inflammatory arthritis (Cosenza et al., 2018).

We evaluated the effect of BMSC-related sEVs on persistent pain in the mouse orofacial pain models. We showed that injection of BMSC-related sEVs attenuated pain hypersensitivity induced by CCI-ION or tendon injury as indicated by decreased mechanical hypersensitivity and reduced aversion to noxious stimulation. The antihyperalgesic effect of sEVs was observed in both female and male animals and the effect was dose-dependent. We previously observed that the pain-attenuating effect of BMSCs appeared to last shorter in female animals (Guo et al., 2016). In the present study, sEVs from naive serum-treated BMSC cultures produced short-lasting antihyperalgesia in male but not female mice, suggesting a subtle sex difference in responsiveness to this therapeutic manipulation. The antihyperalgesia of BMSC sEVs was blocked by GM4869, the antagonist of exosome secretion, suggesting that the effect was not related to other co-isolated soluble mediators and indeed was mediated by MSC-derived exosomes. The use of GM4869 helps to exclude the involvement of ectosomes/microparticles in the observed antihyperalgesia since GW4869 inhibits exosome release but increases microvesicle secretion from the plasma membrane (Menck et al., 2017). Our results support that sEVs/exosomes derived from MSCs could be a surrogate of parent cells for therapeutic outcomes (Ren, 2019). The use of sEVs/exosomes could avoid side effects related to MSC transplantation, such as pulmonary embolism after systemic transplantation (Jung et al., 2013).

The mechanisms underlying the analgesic effect of MSC exosomes are still elusive. MSCs produce antihyperalgesia through immune regulation (Guo et al., 2017). sEVs/exosomes most likely also act on target cells through their cargo molecules with immunomodulatory potential, such as anti-inflammatory cytokines [IL-10, transforming growth factor beta (TGF-β)] (Kordelas et al., 2014; Toh et al., 2018; Ren, 2019) and miRNAs (McDonald et al., 2014). MSC exosomes inhibited the transcription of pro-inflammatory genes, including IL-1β, in the temporomandibular joint (TMJ) osteoarthritis model (Zhang et al., 2019). In the L5/6 spinal nerve ligation model, MSC-exosomes downregulated glia markers glial fibrillary acidic protein (astrocytes) and ionized calcium-binding adaptor molecule 1 (microglia) and pro-inflammatory cytokines TNF-α and IL-1β, and upregulated anti-inflammatory IL-10, associated with a reduction of behavioral hyperalgesia (Shiue et al., 2019; Hsu et al., 2020). Treatment with MSC-derived extracellular vesicles increased immune-suppressive T regulatory cells while attenuating chronic pelvic pain in rats (Peng et al., 2021). The interactions between MSC-exosomes and host cells and their downstream effectors require further investigation.

Interestingly, the *prior injury condition* in which sEVs were isolated favored the pain-relieving effect of sEVs. sEVs isolated from the serum of BMSC-treated injured animals attenuated hyperalgesia for at least 24 h, while sEVs from BMSC-treated naive animals only attenuated hyperalgesia at 3 h after injection. sEVs from the BMSC culture treated with a serum of TL rats were antihyperalgesic, but sEVs from the BMSC culture treated with a serum of naive animals were ineffective. It is rather surprising that we noted this phenomenon, although this is consistent with the previous findings that BMSCs did not affect behavioral pain in non-injured naive animals (Guo et al., 2011). This state-dependent pain relief has been observed in other systems, such as the GluN receptor antagonist that attenuates inflammatory hyperalgesia but does not affect normal nociception (Ren et al., 1992). It has been shown that the injection of exosomes derived from BMSCs not exposed to injury conditions had minimal effect on the mechanical sensitivity of the osteoarthritic rat (He et al., 2020), suggesting the relative ineffectiveness of exosomes derived from non-injured conditions. Multiple injections of exosomes may improve their analgesic efficacy. A single tail vein injection of BMSC exosomes produced about 50% reversal of the vocalization threshold in response to pressure stimulus in a mouse low back pain model (Li et al., 2020). In the rat TMJ osteoarthritis model, however, repeated weekly injections of exosomes from normal human embryonic stem cells fully reversed mechanical hypersensitivity, but note that the first and second injections had no effect, and the maximal effect occurred after the 6th injection (Zhang et al., 2019). Similarly, weekly injections of ECs from MSCs produced antinociception at the late 4-6 weeks but not at week 2 of the injection (Peng et al., 2021). The other way to improve the analgesic effect of exosomes is to slowly deliver exosomes via an alginate scaffold, which produced significant pain relief in nerve-injured rats one day after scaffold implantation (Hsu et al., 2020).

Our results indicate that BMSC-related sEVs produce antihyperalgesia similar to that of BMSCs, and importantly, it is necessary to have prior interactions between BMSCs and injury conditions for producing efficacious pain-relieving sEVs. We have shown that interactions of BMSCs with host immune cells/mediators are critical to their antihyperalgesic effect (Guo et al., 2017). Studies suggest that “*Primed MSCs*” are therapeutically more effective (see Ren, 2019, for a review). MSCs may be altered into a pro- or anti-inflammatory phenotype by differential Toll-like receptor priming (Waterman et al., 2010). MSCs pre-treated with IL-1β produced greater analgesia (Li et al., 2017). Thus, the injury of the host prompts an environment that is targeted by MSCs and their secreted exosomes to produce antihyperalgesia.

The molecular content of MSC exosomes would be dictated by the environment that their parent MSCs encountered. sEVs released *from “primed MSCs”* may possess therapeutic cargo different from those derived from “naive MSCs”. Exosomes derived from macrophage culture subject to LPS stimulation carry an increased number of inflammation-resolving miRNAs (McDonald et al., 2014). The immune inhibitory effect of MSC-derived EVs is enhanced by priming MSCs with inflammatory cytokines (Di Trapani et al., 2016). On the other hand, exosomes derived from interleukin-10 knockout endothelial progenitor cells contain different proteins that are associated with diminished myocardial functional repair (Yue et al., 2020). Thus, the analgesic efficacy of MSC exosomes may be optimized by priming MSCs with inflammatory mediators or specific injury conditions.

EV-based therapy has become an important component of biological medicine that uses cell-derived materials (Lener et al., 2015). The use of EVs/exosomes may get around the side effects associated with cell transplantation. Numerous systemic DiR-labeled MSC exosomes are found in the injured rat spinal cord at 3 and 24 h after injection (Lankford et al., 2018), but most systemically infused MSCs are trapped in the lungs and cannot reach the injured site, and may induce pulmonary embolism (Lee et al., 2009; Jung et al., 2013; Guo et al., 2017). Our results endorse the use of BMSC-derived sEVs/exosomes in chronic pain management and point to the importance of the state of the host for BMSC-exosomes’ action.

## SUPPLEMENTAL INFORMATION

Supplementary information for this article includes 2 figures that can be found with this article online.

## Author Contributions

W.G.,: conception and design, BMSC cultures and exosome isolation, collection and assembly of behavioral, and western blot data, data analysis and interpretation, manuscript writing. J-L.Y.: data collection and analysis. H.K. and K.D.M. BMSC procedures and design. F.W. and K.R. conception and design, data analysis and interpretation, and manuscript writing. All authors gave final approval of the manuscript.

## Acknowledgements

We thank Ms. Shiping Zou for technical assistance. This work was supported by Maryland Stem Cell Foundation grant 2014-MSCRFI-0584; National Institutes of Health grant DE025137.

---

**Supplementary Fig. 1.**
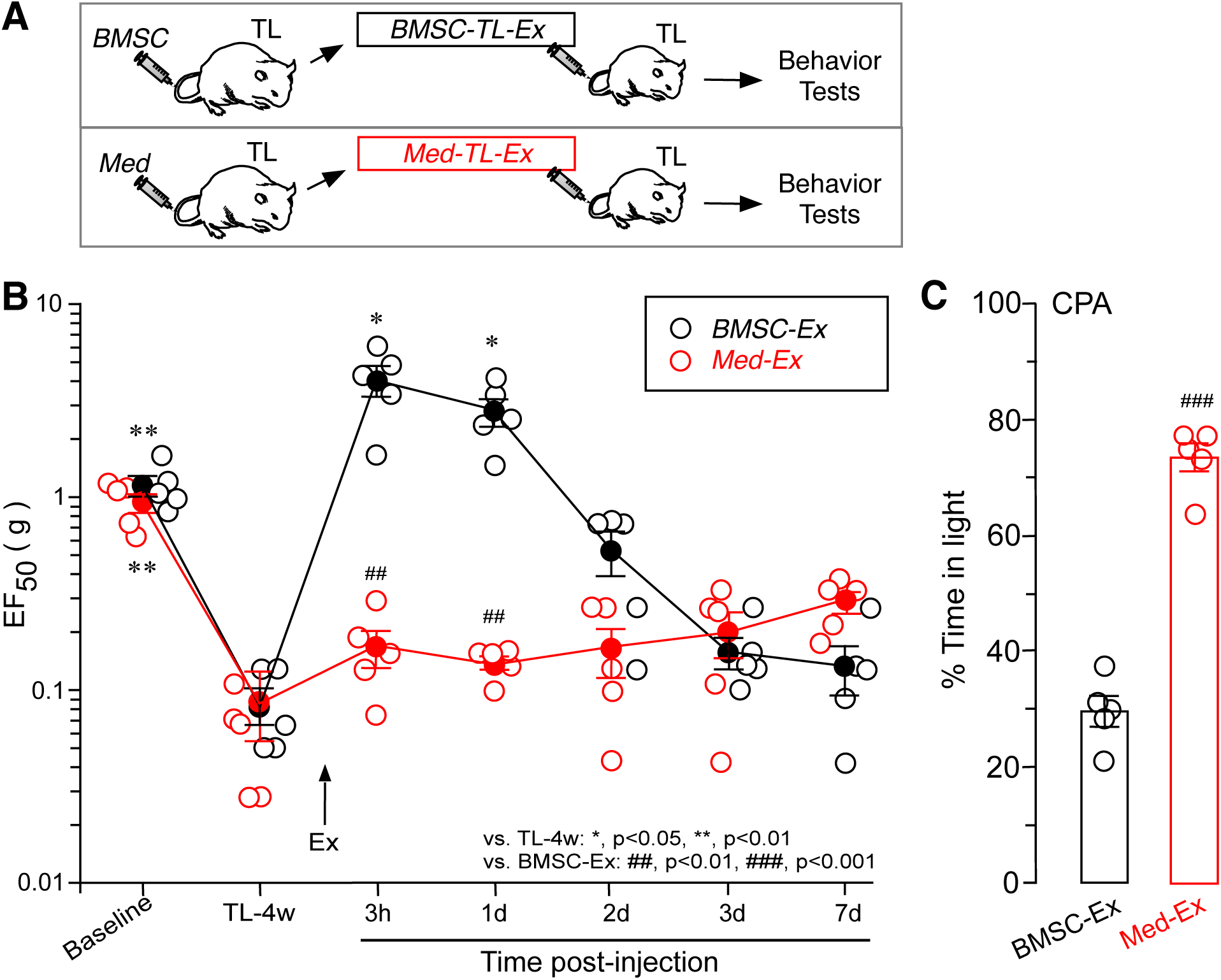
Anti-hyperalgesia by serum-derived sEVs/exosomes in mice after tendon injury. **A**. Flowchart of the experiment. BMSCs or culture medium was injected (i.v.) into the TL rats, and exosomes were isolated from the serum of the donor animals. Isolated exosomes were injected into mice at 4 w after TL, and behavioral tests were conducted at 3 h to 7 d after exosome injection. **B**. An IV injection of exosomes (0.02 ml) isolated from BMSC-treated TL animals (BMSC-TL) elevated EF_50_ at the 3 h and 1 d timepoints, indicating attenuation of behavioral hyperalgesia in TL mice as compared to exosomes from culture medium-treated animals (Med-Ex). **C**. BMSC-Ex reduced %Time in the light chamber in CCI mice at 1 d after injection. *, p<0.05, **, p<0.01, vs. TL-4 w; ###, p<0.001, vs. BMSC-TL. N=5/Group. Post-hoc comparisons after one-way or two-way ANOVA with repeated measures.

**Supplementary Fig. 2.**
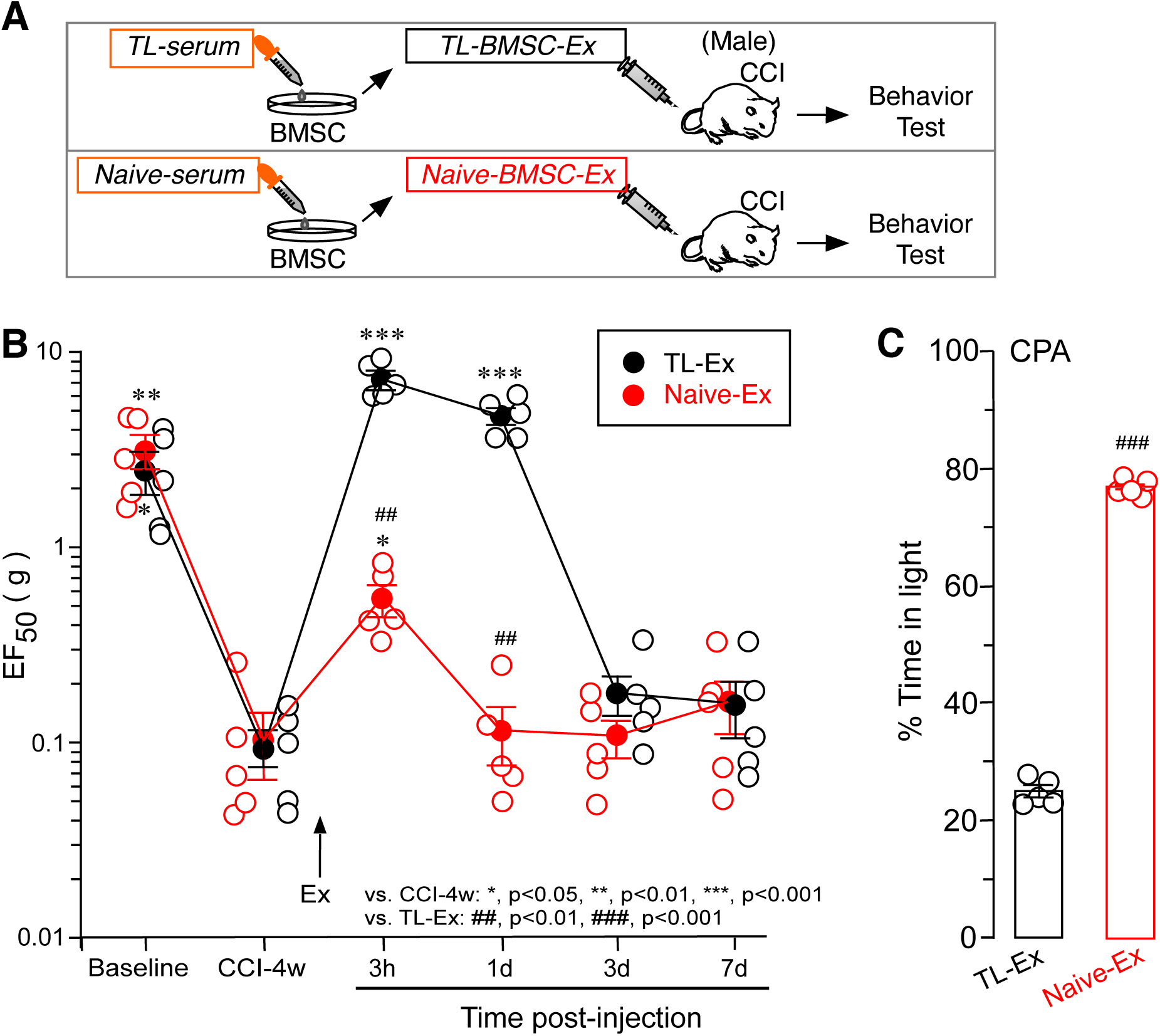
The effect of BMSC culture-derived sEVs/exosomes on behavioral hyperalgesia in male mice. **A**. Flowchart of the experiment. The BMSC culture was primed with serum (20%, 37°C, 16 hr) from TL or naive rats one week after seeding. Ten milliliters of culture medium were used to isolate exosomes. **B**. An IV injection of exosomes (0.02 ml) isolated from the BMSC culture primed with serum from TL animals (TL-Ex) elevated EF_50_ in CCI mice, compared to exosomes from the BMSC culture treated with naive serum (Naïve-Ex). **C**. TL-Ex reduced % Time in the light chamber in CCI mice at 1d after injection, suggesting a reduced aversion to a noxious stimulus. *, p<0.05, **, p<0.01, ***, p<0.001, vs. CCI-4w; ##, p<0.01, ###, p<0.001, vs. TL-Ex 1x. N=5/Group. Post-hoc comparisons after one-way or two-way ANOVA with repeated measures.

